# Somatosensory cortical representations of assimilation effect by vibrotactile stimulation

**DOI:** 10.1101/2023.08.27.555001

**Authors:** Ji-Hyun Kim, Dooyoung Jung, Junsuk Kim, Sung-Phil Kim

**Affiliations:** Department of Biomedical Engineering, Ulsan National Institute of Science and Technology, Ulsan, South Korea; School of Information Convergence, Kwangwoon University, Seoul, South Korea

**Author notes:** **Corresponding author 1:** Department of Biomedical Engineering, Ulsan National Institute of Science and Technology, 50 Unist-gil, Eonyang-eup, Ulsan 44191, South Korea ***Email:** (S.-P. Kim). **Corresponding author 2:** School of Information Convergence, Kwangwoon University, 20 Kwangwoon-ro, Nowon-gu, Seoul 01897, South Korea ***Email:** (J. Kim).

**Keywords:** Vibrotactile, Somatosensory cortex, fMRI, Assimilation effect, Connectivity

## Abstract

This study investigated neural activity related to the assimilation effect in the perception of vibrotactile stimuli. The assimilation effect refers to a tactile perceptual bias in which the perception of the vibrotactile frequency on one finger is biased towards a distracting vibrotactile stimulus targeting other types of mechanoreceptors on the different finger. The assimilation effect occurs not only between fingers on the same hand (in-hand) but also between fingers on different hands (across-hand). These behavioral aspects of the assimilation effect lead to an assumption that neural processes related to the assimilation effect would involve the integration of different tactile information mediated by the somatosensory cortex. We addressed this hypothesis by investigating brain responses using functional magnetic resonance imaging (fMRI) to vibrotactile stimuli that induced assimilation effect under in-hand and across-hand conditions. We first observed that the vibrotactile stimuli presented in this study activated primary (S1) and secondary (S2) somatosensory cortices. Yet, neural responses in these regions did not show correlations with individual assimilation effects, indicating that neural processing of vibrotactile signals in S1 and S2 would not be directly linked to the assimilation effect. Instead, we found that connectivity between S1 and medial prefrontal cortex (mPFC) was correlated with individual in-hand assimilation effects and that connectivity between S2 and posterior parietal cortex (PPC) was correlated with individual across-hand assimilation effects. These results suggest that the assimilation effect may be related to tactile information integration via functional connections between the somatosensory cortex and higher-order brain regions.

## 1. Introduction

The somatosensory nervous system processes tactile information from various mechanoreceptors distributed over the body (Abraira and Ginty, 2013; Johansson and Flanagan, 2009). Different types of tactile stimuli are separately delivered by individual tactile afferents connected to cutaneous receptors (Johnson, 2001) that are broadly divided into rapidly adapting (RA) and slowly adapting (SA) receptors (Johansson and Vallbo, 1983). Non-human primate studies have shown that neural signals delivered by RA and SA afferents arrive at distinct columns and layers of area 3b in the primary somatosensory cortex (S1) (Mountcastle et al., 1969; Sur et al., 1984) and that S1 neurons receiving either SA or RA afferent input form distinct clusters (Chen et al., 2001; Friedman et al., 2004).

Although it remains unknown how S1 neurons in humans are distributed according to SA and RA afferent input, findings in human behavioral studies have indicated that different types of tactile stimuli are perceived separately. A behavioral study reported that adaptation to flutter, which predominantly causes desensitization of RA type I (RA-I) and SA type I (SA-I) afferents, did not affect fine-texture discrimination mediated by RA type II (RA-II) afferent (Bensmaïa and Hollins, 2005). It suggests that the human somatosensory cortex may also represent different types of tactile stimuli in a segregated fashion.

However, even though individual types and locations of tactile stimuli would be perceived independently, diverse tactile information received concurrently needs to be integrated in the brain to enable coherent tactile perception, such as texture recognition (Rostamian et al., 2022). One of those perceptual phenomena can be found in a biased perception known as the assimilation effect (Sherif et al., 1958). A behavioral study has demonstrated the assimilation effect in vibrotactile perception where the judgment of the frequency of a target vibrotactile stimulus given to one finger shifted towards the frequency of a distractor vibrotactile stimulus concurrently given to the adjacent finger or the same finger in the opposite hand (Kuroki et al., 2017). Research on this vibrotactile assimilation effect can shed a light on lesser-known mechanisms underlying the integration of various types of tactile information. However, neural correlates of the assimilation effect in tactile perception are still unclear.

Therefore, this study aims to investigate neural activity related to the assimilation effect in vibrotactile perception via functional magnetic resonance imaging (fMRI) of the human brain. In the human EEG experiments using visual luminance stimuli, responses of occipital and parietal cortices were observed when the assimilation effect occurred (Acaster et al., 2021). A neuroimaging study also showed that neural responses in the secondary somatosensory cortex (S2) were increased in subjects who exhibited the assimilation effect when perceiving flavor stimuli (Davidenko et al., 2018). Following the previous assimilation effect studies with different sensory modalities, we assume that the parietal cortex would be related to the assimilation effect. Especially, we assume that S2, involved in the high-level processing of tactile information delivered from S1 and the thalamus (Eickhoff et al., 2007), may merge different types of tactile signals, likely resulting in interference between tactile signals.

We also assume that the assimilation effect can influence connectivity between the somatosensory cortex and other brain regions, which would be involved in higher-order tactile processing, such as tactile discrimination (Hartmann et al., 2008). We employ the same experimental design as in the previous behavioral study to test the assimilation effect, in which subjects discriminate the frequencies of vibrotactile stimuli activating RA-I afferents while simultaneously receiving a distractor vibrotactile stimulus activating RA-II on the same hand or different hands (Kuroki et al., 2017). This design includes a control condition where a distracting RA-II stimulus is replaced by RA-I. As such, changes in regional activation or inter-regional connectivity by the assimilation effect compared to the control condition can merely reflect neural responses to multiple types of vibrotactile stimuli. To address this confounding, we further assume that neural substrates of assimilation effects would exhibit correlations with individual perceptual assimilation effects.

## 2. Materials and Methods

### 2.1. Subjects

Thirty-nine subjects (20 females; mean age 25.4 years old; age range 19-35 years old) with no contraindications against MRI and no history of neurological disorders participated in this study. Only right-handed subjects were recruited to control the handedness effect. Each subject participated in the pre-experiment, behavioral experiment, and fMRI experiment. Seven subjects whose behavioral data could not be fitted by a general psychometric curve were excluded during the processing of behavioral data (see below for details in *2.6*. *Behavioral data analysis*). Other six subjects were also excluded during fMRI data analysis, three of whom were excluded because of their head movements and the other three because of the lack of individual region of interests (ROIs) (see below for details in *2.9*. *Determination of local region of interests (ROI)*). Consequently, the data of twenty-six subjects were used in the main analyses. This study was approved by the ethics committee of the Ulsan National Institute of Science and Technology (UNISTIRB-21-54-C). The study was conducted according to the Declaration of Helsinki. All subjects were informed of the study objectives and experimental procedures and voluntarily submitted a written consent form.

### 2.2. Vibrotactile stimuli

We adopted the design of vibrotactile stimuli from the previous study, with which the assimilation effect on tactile perception was demonstrated (Kuroki et al., 2017). We initially set two primary frequencies of vibrotactile stimuli: 1) a target frequency of 30 Hz to predominantly activate RA-I afferents; and 2) a distractor frequency of 200 Hz to predominantly activate RA-II afferents. Note that we used a 200-Hz stimulus, whereas the study by Kuroki *et al*. used a 240-Hz stimulus because the piezoelectric tactile stimulator (PTS) used in this study elicited a resonance effect for stimuli over 200 Hz, significantly reducing the stimulus amplitude. In addition, the vibrotactile frequency of 200 Hz is sufficiently high to stimulate RA-II receptors (Dykes et al., 1981). Then, we set the frequency of six comparison vibrotactile stimuli around the target frequency to assess the perception of vibrotactile stimulation as 15, 21, 25, 36, 42, and 60 Hz. We synthesized all the vibrotactile stimuli with the sinusoidal signals using MATLAB 2019b (Mathworks, Inc. Natick, MA, USA). In addition, Tukey windowing was used to reduce the prominence of skin deformation on the onset and offset of stimulation (Kuroki et al., 2017). We used an MRI-compatible PTS equipped (6-mm diameter) to provide vibrotactile stimulations on the skin (Dancerdesign, St. Helens, UK). Two PTSs were attached to the middle and index fingertips of the left hand and one to the index fingertip of the right hand. The amplitude of the target stimulus (30 Hz) was fixed at 0.4 arbitrary unit (AU). Then, the distractor and comparison stimuli amplitudes were adaptively determined for each subject through the pre-experiment.

### 2.3. Pre-experiment

The pre-experiment was designed to balance the perceived intensities across all the vibrotactile stimuli with different frequencies in each subject. This procedure was required because the PTS used in this study increased the amplitude of generated sinusoidal signals as the frequency increased. Also, it is known that perceived vibrotactile intensity varies with the frequency of vibrotactile stimuli (Prsa et al., 2021). Therefore, we aimed to ensure that each subject perceived the same intensity for all the vibrotactile stimuli by adjusting the amplitude of every stimulus for each subject. To set the initial values of the amplitude of each stimulus, we utilized a dataset generated by our laboratory from a different study where fifty-seven subjects performed the same task to discriminate vibrotactile stimuli provided by the same apparatus used in this study (Jeong et al., 2022). We applied the staircase method to determine the amplitude of each stimulus so that the perceived intensity of every stimulus was equalized in each subject (Wetherill and Levitt, 1965). Then, we averaged the amplitudes across all fifty-seven subjects per stimulus and used them as initial values for the current study (Fig. 1).

In the pre-experiment, subjects sat on a chair with both hands extended forward, facing their palms downward. Subjects were instructed to gaze at the monitor displaying a black cross on a grey background during the pre-experiment. The black cross changed to white to cue the start of a trial, and a pair of vibrotactile stimuli were simultaneously delivered to the index and middle fingers of the left hand for 1 s. One of the pair was the target stimulus (30 Hz) with a fixed amplitude (0.4). The other was either one of the comparison stimuli (15, 21, 25, 36, 42, and 60 Hz) or the distractor stimulus (200 Hz) with an adaptive amplitude (Fig. 2A, B). The fingers stimulated by the target stimulus and the comparison/distractor stimulus were fully randomized. After a 1-s stimulation period, the word “response” was displayed on the screen, and subjects responded which stimulus intensity was higher. Since their hands were fixed, subjects responded by moving their foot. Subjects pressed the left foot pad if the stimulus to the middle finger felt stronger and the right foot pad if the stimulus to the index finger felt stronger. A trial ended with the subjects’ response, and a subsequent trial began after a 2-s inter-trial interval. We presented a stimulus pair for 30 trials in a single run for each comparison or distractor stimulus. If subjects rated one of the stimuli stronger in more than 19 trials, we determined that the perceived intensities of the two stimuli did not match with each other and adjusted the amplitude of the comparison/distractor stimulus using the 3-up/1-down and 3-down/1-up staircase algorithm (Wetherill and Levitt, 1965). We repeated the runs until the number of trials subjects rated one of the stimuli stronger did not exceed 19. We repeated this procedure for every comparison/distractor stimulus.

**Fig. 1.**
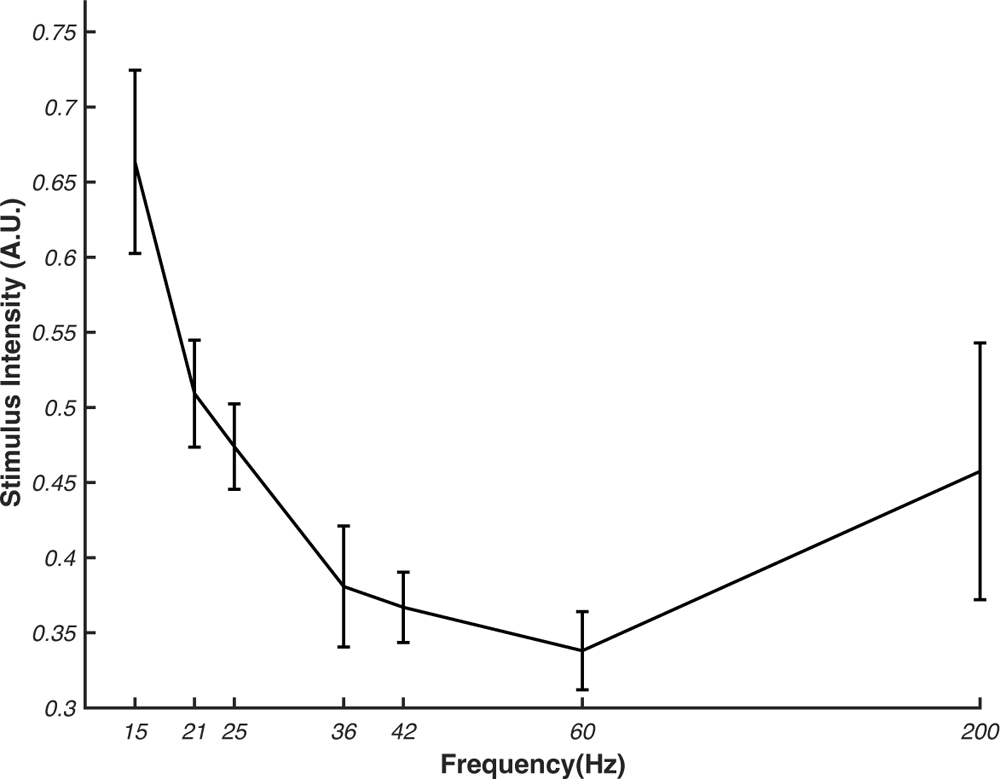
Mean adjusted amplitudes of vibrotactile stimuli. The amplitudes of the non-target stimuli are adjusted using the 3-up/1-down and 3-down/1-up algorithms based on the subjects’ responses. The amplitude of each non-target stimulus is adjusted to match its perceptual intensity to that of the target stimulus (at 30 Hz). The amplitude of the target stimulus is set to 0.4. The error bars denote the standard error of mean across the subjects (N = 57).

**Fig. 2.**
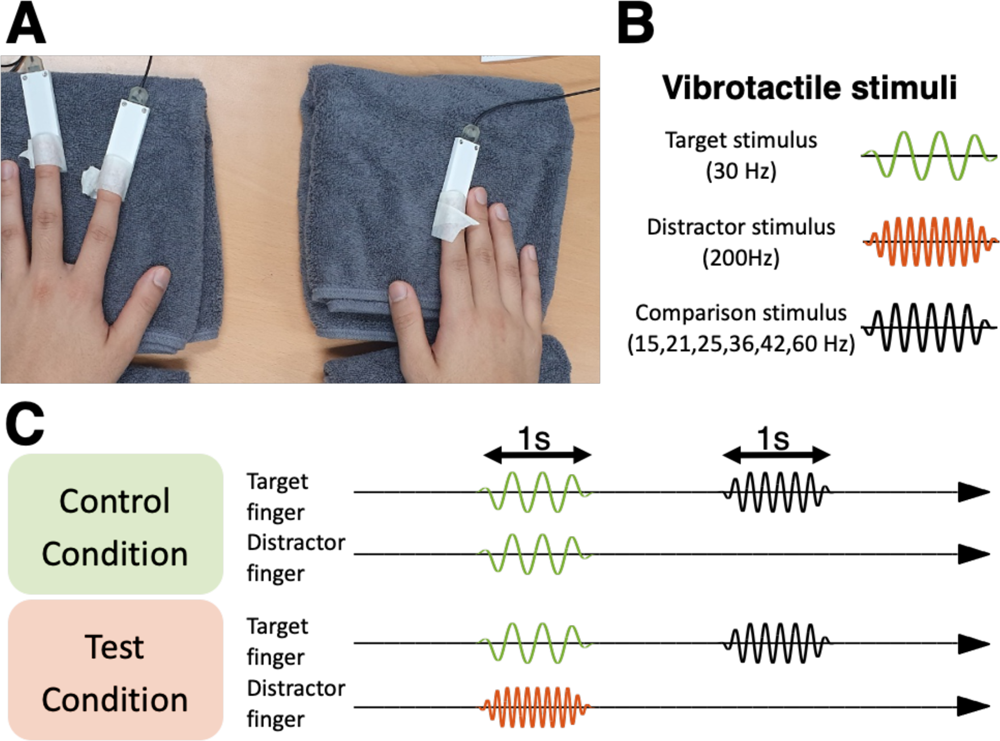
Experimental setup and stimuli. (A) Attachment of stimulators to the fingers for every experimental condition. The subject conducts the experiment with placing the hands on a towel. Three stimulators are attached to the left middle fingertip, left index fingertip, and right index fingertip. (B) Vibrotactile stimuli used in the study are divided into three types: target stimulus at 30 Hz, distractor stimulus at 200 Hz, and comparison stimuli at 15, 21, 25, 35, 42, and 60 Hz. (C) The conditions of stimulus presentation. Under the control condition, the target stimulus is delivered to both the target and distractor fingers simultaneously for 1s. After a 1s inter-stimulus interval, one of the comparison stimuli is delivered to the target finger for 1 s. Under the test condition, the target stimulus is delivered to the target finger and the distractor stimulus is delivered to the distractor finger at the same time for 1s. After a 1s inter-stimulus interval, one of the comparison stimuli is delivered to the target finger for 1 s.

### 2.4. Behavioral experiment

Subjects performed the behavioral experiment in the same room after the pre-experiment. The break time between the pre-, and behavioral experiments was less than 5 min. We adopted the behavioral experimental paradigm described in the previous study (Kuroki et al., 2017). The behavioral experiment proceeded under three conditions that differed in terms of how vibrotactile stimuli were given: Non-Distractor (ND), Across-Finger (AF), and Across-Hand (AH) conditions. However, in all conditions, the experiment underwent the same sequence of three periods: 1) a target period with the presentation of the target stimulus (30 Hz) (1 s); 2) an interstimulus interval (ISI) (1 s); and 3) a comparison period with the presentation of one of the comparison stimuli (1 s).

In the ND condition, the target stimulus was given to a target finger without the distractor stimulus. The target finger was either the index or middle finger of the left hand. In the AF condition, the left index or middle finger different from the target finger (termed as a distractor finger here), was simultaneously stimulated during the target period. The AF condition was further divided into two sub-conditions. The target stimulus (30 Hz) was given to the distractor finger under an AF control condition (AFc), whereas the distractor stimulus (200 Hz) was given under an AF test condition (AFt) (Fig. 2C). The AH condition was the same as the AF condition, except that the target and distractor fingers were assigned as the index fingers of both hands. A trial consisted of vibrotactile stimulations over three periods as described above as well as behavioral responses. We blocked a set of trials according to finger arrangement and task condition. The locations of the target and distractor fingers were alternated in two different finger arrangements for each condition. At the beginning of a block, a text instructing to pay attention to stimulation on the target finger appeared on the screen. Afterward, a black fixation cross appeared on the screen for 1s and turned to white to cue the start of the stimulus presentation. The cross became white only during stimulations and remained black otherwise. After the comparison period, subjects responded which of the target and comparison stimuli vibrated with a higher frequency. Subjects pressed the left foot pad if the frequency of the target stimulus felt higher and the right foot pad that of the comparison stimulus felt higher.

Each of the six comparison stimuli was presented 10 times in a random order within a block so that every block contained 60 trials. Subjects performed a total of 10 blocks (600 trials) with 2 blocks under the ND (the left index or middle finger was target in each block), 4 blocks under the AF (the left index or middle finger was target in each block two times, respectively), and 4 blocks under the AH conditions (the left index or right index finger was target in each block two times, respectively) (Fig. 3A). In a single block under the AF or AH condition, control (AFc or AHc) and test (AFt or AHt) conditions were mixed in a pseudorandom order to balance the number of presentations between the comparison stimuli. The order of ND, AF, and AH blocks was fully randomized.

**Fig. 3.**
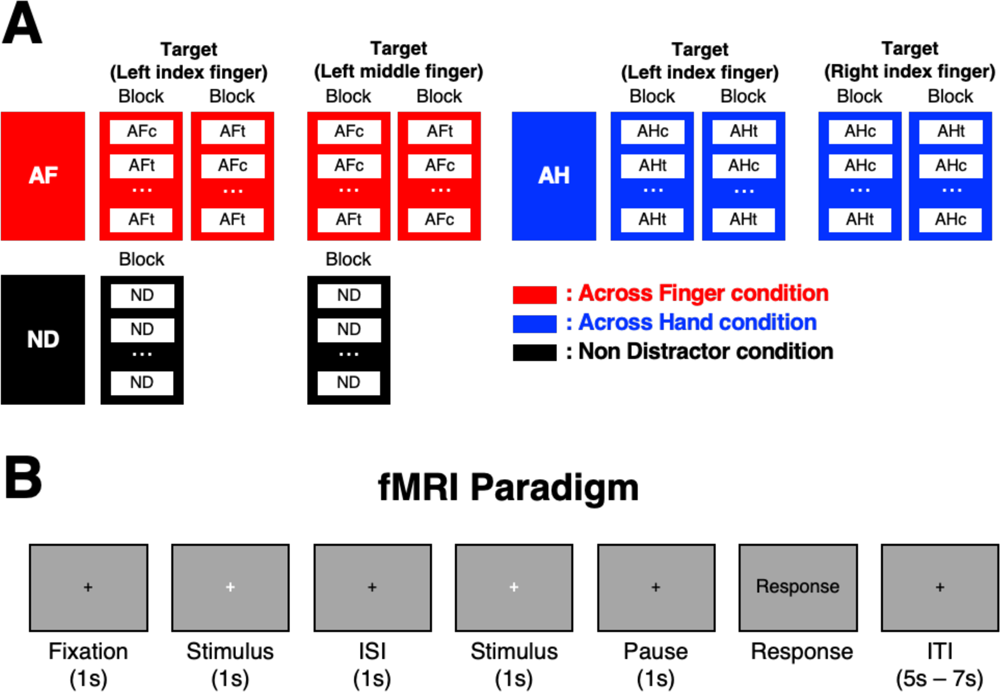
Experimental paradigm. (A) Conditions of the behavioral task. The subject performs the behavioral task by discriminating higher frequency between the target and comparison stimuli in each trial under three conditions: across-finger (AF), across-hand (AH), and non-distractor (ND). In the AF and AH conditions, stimuli are presented in either the control condition (AFc, AHc) or the test condition (AFt, AHt) for each trial. The control and test conditions are randomly presented in a block of the behavioral condition. Each block contains 30 trials of the control and 30 trials of the test conditions. Two blocks are included in each finger configuration: left index finger as target or left middle finger as target in the AF condition; and left index finger as target and right index finger as target in the AH condition. There is no distractor stimulus in the ND condition. (B) Experimental design for fMRI experiment. A white cross appears at the center of the screen when stimuli are presented. The subject responds higher frequency when the word “response” is displayed. An inter-trial interval (ITI) is randomly determined between 5s and 7s.

### 2.5. fMRI experiment

The fMRI experiment was conducted 2.15±1.29 days on average after the behavioral experiment. Three PTSs were attached to the fingers in the same way as in the behavioral experiment while the subjects’ palms were placed in a downward position. The fMRI experimental paradigm was identical to the behavioral experimental paradigm except that each comparison stimulus was presented four times (instead of ten) per block due to time limitation of MR scanning. Subjects responded by pressing the foot pad in the MR scanner, similar to the behavioral experiment (Fig. 3A). Also, an inter-trial interval was set to 5 to 7s in order to meet the requirement for the hemodynamic response function (HRF) to return to baseline (Fig. 3B).

### 2.6. Behavioral data analysis

We estimated the psychometric function of behavioral responses in the perceptual discrimination of vibrotactile frequency using the Palamedes toolbox 1.10.11 for MATLAB (Prins et al., 2018). The behavioral data collected from both the behavioral and fMRI experiments were aggregated and used for the estimation. For each comparison stimulus under each condition in each subject, we calculated a ratio of the number of trials where the frequency of the comparison stimulus was perceived higher than that of the target stimulus to the total number of trials the comparison stimulus was given. There were 28 trials – 20 from behavioral and 8 from fMRI experiments – for each of ND, AFc, AFt, AHc and AHt condition. The logistic function was used as the psychometric function and fitted to the calculated ratio of each comparison stimulus under each condition in each subject. The Nelder-Mead simple search algorithm was used to maximize the likelihood of psychometric function (Nelder and Mead, 1965). The fitted psychometric functions was deemed to be abnormal when the ratio constant or decrease as the frequency of the comparison stimulus increased. Seven subjects who did not show a normal psychometric function for any of the five conditions above were excluded from the analysis.

From the psychometric functions for each subject, we calculated the point of subject equality (PSE), which was the estimated frequency of the comparison stimuli corresponding to 50% of the ratio on the fitted psychometric curve. The previous behavioral study revealed that the PSE difference between the test and control conditions could estimate the degree of the assimilation effect (Kuroki et al., 2017). The significance of the PSE difference between AFc, ND, and AFt as well as between AHc, ND, and AHt was statistically evaluated using the Wilcoxon signed rank test (α = 0.05).

### 2.7. MRI data acquisition and preprocessing

The MRI scanning was conducted using a 3 T scanner (Magnetom TrioTim, Siemens, Germany) with a 64-channel head coil at the Center for Neuroscience Imaging Research (CNIR) in Suwon, Republic of Korea. 3-D functional images were acquired using a slice-accelerated multiband gradient-echo-based echo planar imaging (EPI) sequence using T2*-weighted blood oxygenation level-dependent (BOLD) contrast, covering the whole brain region (multiband acceleration factor = 2, number of slices = 72, repetition time (TR) = 2,000 ms, echo time (TE) = 35 ms, flip angle = 90°, Field of view (FOV) = 200 mm, slice thickness = 2 mm and voxel size = 2.0 × 2.0 × 2.0 mm^3^). Also, anatomical high-resolution images were obtained (T1-weighted 3D MPRAGE sequence, TR = 2,300 ms, TE = 2.28 ms, flip angle = 8°, FOV = 256 mm, and voxel size = 1.0 × 1.0 × 1.0 mm^3^). Functional images were preprocessed using SPM12 software (Wellcome Department of Imaging Neuroscience; London, UK). The preprocessing of functional images underwent a series of steps, including slice-timing correction, realignment, co-registration, segmentation, spatial normalization to the Montreal Neurological Institute (MNI) template, and smoothing with a 6-mm full-width-half-maximum isotropic Gaussian kernel. Three subjects were excluded during preprocessing because their head movements exceeded the voxel size (2 mm).

### 2.8. Univariate analysis of fMRI data

Preprocessed fMRI data were analyzed using the general linear model (GLM) to find individual voxels activated by the vibrotactile stimulation. The BOLD responses of single voxels were modeled using the SPM canonical hemodynamic response function (HRF).

Regressors of the design matrix in the GLM analysis within a single block included: 1) the onset of the target period under the control condition; 2) the onset of the target period under the test condition; 3) the onset of the comparison period regardless of the condition; and 4) the onset of subjects’ response. Additional regressors of no interest for motion correction parameters and a linear scanner drift were included in the design matrix. We created a contrast by beta estimates for the stimuli condition to determine voxels activated by vibrotactile stimuli. Then, the group-level analysis was conducted with a one-sample t-test using the contrast images obtained from the contrast analysis above. The resulting contrast maps were thresholded to correct for multiple comparisons using cluster-level inference (Woo et al., 2014). We used a significance threshold of p<0.05 family-wise error (FWE) corrected for multiple comparisons at the cluster level, with a height threshold at the voxel level of p<0.001.

### 2.9. Determination of local region of interests (ROI)

After obtaining the contrast maps, we determined the local region of interests (ROIs) in each subject for each of the AF and AH conditions by assuming that each subject would have a different peak activation location even in the same anatomical region. ROIs were determined based on the following criteria. First, we restricted the ROIs to the somatosensory cortex based on the stimuli vs baseline contrast since the primary aim of the study was to identify neural correlates of the assimilation effect within this region. Second, we sought individual peak activation locations within a volume with a 12-mm diameter centered at the local maximum determined by the group analysis. Third, an ROI for each subject was determined as a set of voxels that showed the same contrast (stimuli vs. baseline, threshold of p<0.05, uncorrected) and were located within an 8-mm distance from the individual peak activation location. Note that we used stimuli vs baseline contrast to determine ROIs as we aimed to find local somatosensory cortical regions involved in processing a set of tactile stimuli presented in this study. Based on ROIs determined using stimuli vs baseline contrast, we further analyzed whether neural activities in those ROIs were correlated with the behavioral outcomes of the assimilation effect. We excluded the data of three subjects from the analyses using ROIs because no ROI satisfying the selection criteria was found in these subjects.

### 2.10. Percent signal change analysis of fMRI data

We first analyzed a relationship between local regional activity and the behavioral outcomes of the assimilation effect. We investigated whether the percent signal change (PSC) in individual local ROIs over the somatosensory cortex was related to individual PSE values. Using the Marsbar toolbox (Brett et al., 2016), we first calculated the PSC of each local ROI under each of the AFt, AFc, AHt, and AHc conditions by multiplying the beta weight for the condition by the peak value of the single-trial regressors and dividing it by the beta weight for the constant regressor. Next, we examined whether the PSC in each ROI was statistically different between the test and control conditions (i.e., AFt vs AFc and Aht vs Ahc) using the Wilcoxon signed rank test (α = 0.05). Differences in the PSC between the test and control conditions do not necessarily support that the corresponding neural responses represent the assimilation effect because those differences could simply reflect differential neural responses to multiple tactile stimuli (RA-I and RA-II) compared to a single stimulus (RA-I). Thus, we further investigated whether the PSC differences were correlated with the degree of behavioral change by the assimilation effect. To this end, we calculated the Pearson correlation coefficient between the PSC difference and the PSE difference across individual subjects for each ROI, where the PSE difference was measured between the test and control conditions.

### 2.11. Brain connectivity analysis of fMRI data

We analyzed connectivity between somatosensory cortical regions and other brain regions in relation to the behavioral outcomes of the assimilation effect. We assessed connectivity from each ROI to other voxels over the whole brain using the psycho-physiological interaction analysis (PPI) (Friston et al., 1997; O’Reilly et al., 2012). Especially, we used a generalized PPI analysis (gPPI) to improve sensitivity and specificity (McLaren et al., 2012). In the gPPI analysis, we extracted the first eigenvariate BOLD signal from each ROI and deconvolved it using the canonical HRF to obtain an approximation of neural activity. Next, we centered the resulting time series of neural activity and multiplied it with the psychological factors defined as {Test vs Baseline} and {Control vs Baseline}. This psychologically factored neural activity was termed as interaction time series. Finally, we convolved the interaction time series with the canonical HRF again, generating a hemodynamic-level interaction variable (called a PPI term). We reconstructed an individual-level design matrix by adding regressors of the PPI terms as well as the extracted time series of the local ROI to other existing regressors. Then, we applied a GLM to predict the BOLD signals of a target voxel from this design matrix. We computed the contrast of the PPI regressors for {Test vs Baseline} and {Control vs Baseline} using the paired t-test in each subject and utilized the obtained contrast images for subsequent group analysis. To evaluate statistical significance, we set the height threshold at p = 0.001 (uncorrected) and corrected the extent threshold of activation to p < 0.05, accounting for multiple comparisons across the whole brain using FWE correction. After finding brain regions showing significant inter-regional connectivity from somatosensory cortical ROIs by the gPPI analysis, we assessed its relationship with individual behavioral assimilation effects. Similar to the PSC analysis above, we used individual differences in the PSE between the test and control conditions. We defined individual connectivity as the mean of the beta estimates of the gPPI in the identified brain regions. We calculated the Pearson correlation coefficient between individual connectivity and the PSE difference across subjects.

## 3. Results

### 3.1. Behavioral results

We observed the behavioral outcomes reflecting the assimilation effect in both the AF and AH conditions, as in the previous study (Kuroki et al., 2017). Basically, we confirmed that an increase in the frequency of the comparison stimuli (comparison frequency) led to a higher proportion of judgments that the comparison frequency was higher than that of the target stimulus (target frequency). The assimilation effect was revealed by the observation that the psychometric function in the AFt (AHt) condition was shifted towards higher frequency relative to that in the ND and AFc (AHc) conditions (Fig. 4A, B). The PSE was significantly higher in the AFt condition (33.22±5.74 Hz, MEAN±SEM) than ND (29.04±3.41 Hz, p=0.0045) and the AFc (28.39±4.41 Hz, p = 0.0032) conditions, indicating that the perceived target frequency was shifted towards a higher frequency due to the presence of the distractor stimulus (200 Hz) (Fig. 4C). Likewise, the PSE was higher in the AHt condition (32.85±4.16 Hz) than ND (29.04±3.41 Hz, p=0.001) and AHc (29.01±4.20 Hz, p = 0.0012) conditions (Fig. 4D). These results showed that the assimilation effect occurred within the hand as well as across the hands. Although the PSE shift in the AHt condition was slightly smaller compared to that in the AFt condition, a difference in the PSE between the two conditions was not significant (Wilcoxon signed rank test, p = 0.30).

**Fig. 4.**
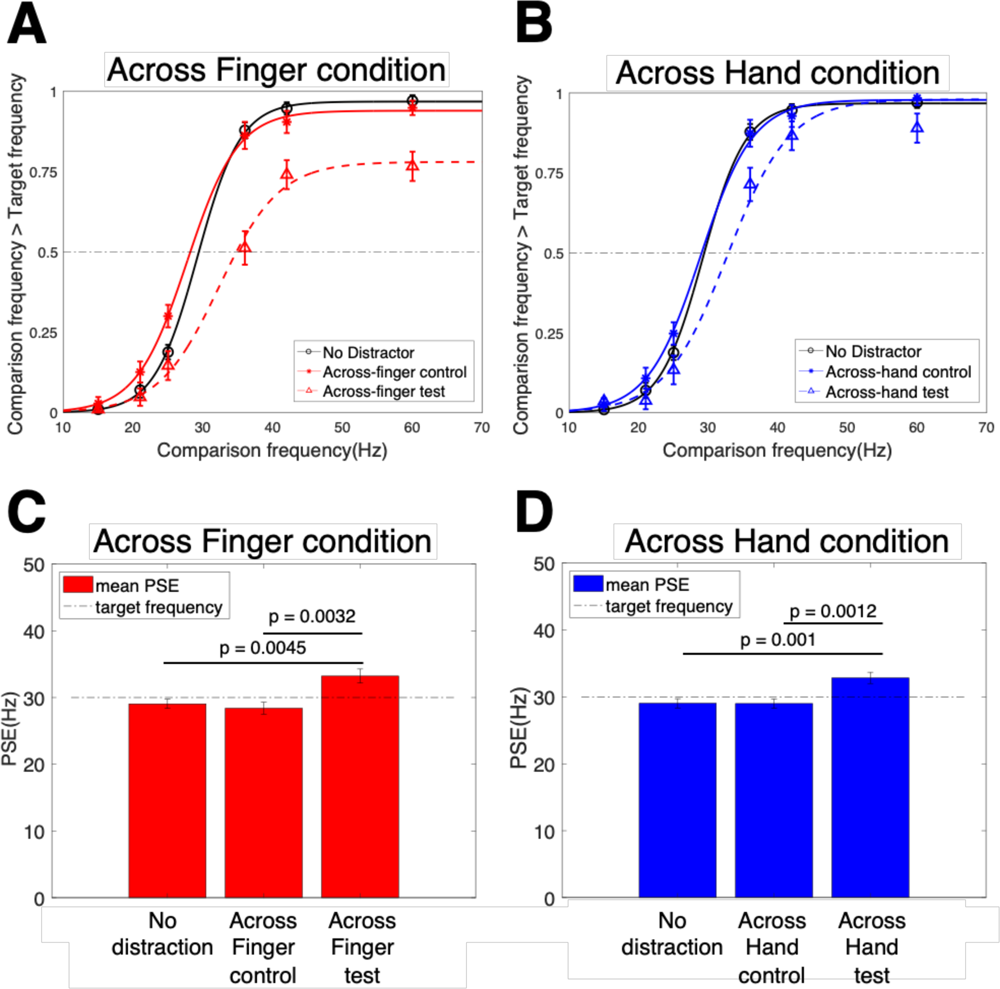
Behavioral results of assimilation effects. (A) Psychometric functions of across-finger (AF) conditions. The mean proportion of trials when subjects judged the comparison stimulus vibrated at a higher frequency than the target stimulus (30Hz) was marked on the vertical axis for six different comparison stimuli at 15, 21, 25, 35, 42, and 60 Hz (N= 26 subjects). The psychometric functions of the non-distractor (ND, black open circle) and the control (AFc, red circle) conditions are similar, while the psychometric function of the test (AFt, red triangle) condition is deviant. (B) Psychometric functions of across-hand (AH) conditions. The psychometric functions of the ND (black open circle) and the control (AHc, red circle) conditions are similar, while the psychometric function of the test (AHt, red triangle) condition is deviant. C. The mean points of subject equality (PSEs) in the AF conditions. The mean PSE in the AFt condition is significantly different from that in the ND (p=0.0045) and that in the AFc condition (Wilcoxon signed rank test, p = 0.0032). There is no difference between the ND and AFc conditions (p = 0.1718) D. The mean PSEs in the AH conditions. The mean PSE in the AHt condition is significantly different from that in the ND (p=0.001) and that in the AHc condition (p = 0.0012). There is no difference between the ND and AFc conditions (p = 0.9274). Error bars indicate the standard error of the mean.

### 3.2. Regional neural activations

The contrast analysis of stimuli versus baseline showed that the vibrotactile stimuli activated the somatosensory cortical regions. In the AF condition, the bilateral secondary somatosensory cortex (S2) and contralateral S1 showed significant activations by the vibrotactile stimulation in the target period (FWE corrected p<0.05, cluster size >10, cluster defining threshold p<0.001) (Fig. 5A). Additional significant responses were observed in other regions including the precentral gyrus, occipital lobe, supplementary motor area, cerebellum, insula, and inferior frontal lobe (FWE corrected p<0.05, cluster size >10, cluster defining threshold p<0.001) (Table. 1). In the AH condition, bilateral S1 and bilateral S2 showed significant activations by the vibrotactile stimulation in the target period (FWE corrected p<0.05, cluster size >10, cluster defining threshold p<0.001) (Fig. 5B). In addition, occipital lobe and left precentral gyrus were also activated (FWE corrected p<0.05, cluster size >10, cluster defining threshold p<0.001) (Table. 1). Bilateral S2 showed significant activations during the test versus control conditions in both AH and AF conditions (Table. 1).

**Fig. 5.**
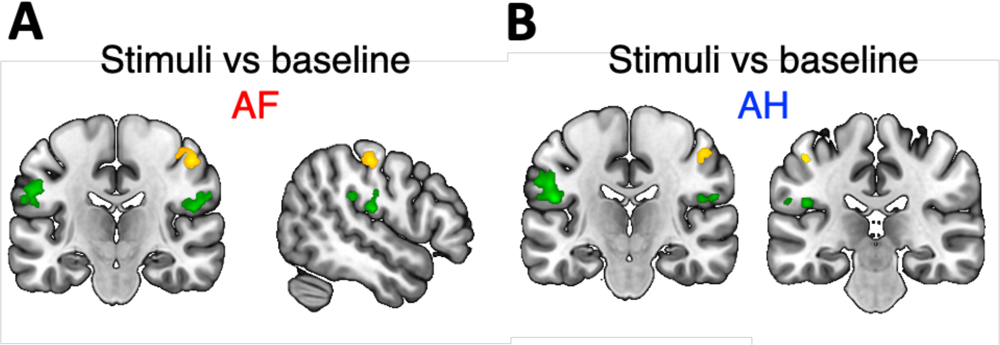
Brain regions activated by vibrotactile stimuli in contrast to baseline. (A) Brain response maps showing brain regions activated by all the vibrotactile stimuli used in this study in the across-finger (AF) condition, resulting from the contrast analysis of stimuli versus baseline. Right S1 contralateral to stimuli (yellow) and bilateral S2 (green) regions show significant responses. (B) Brain response maps of stimuli vs baseline in the across-hand (AH) condition. Bilateral S1 (yellow) and bilateral S2 (green) regions show significant responses. Significance of activations are corrected for multiple comparisons p<0.05 at the cluster level, with a height threshold at the voxel level of p<.001.

**Table 1.** Contrast analysis results. The results from the group-level contrast analysis of stimuli versus baseline are summarized. The 3D coordinates and t-values of each cluster peak point are listed. Significance of activations is corrected for multiple comparisons p<0.05 at the cluster level, with a height threshold at the voxel level of p<.001.

| AF (Stimuli vs Baseline) |  |  |  |  |
| --- | --- | --- | --- | --- |
| Brain region | Coordinates |  |  | T |
|  | x | y | z |  |
| R_S1 | 50 | -20 | 50 | 7.94 |
|  | 44 | -26 | 56 | 4.84 |
|  | 40 | -30 | 48 | 4.71 |
| L_M1 | -56 | 6 | 32 | 6.49 |
|  | -58 | 4 | 22 | 5.87 |
|  | -46 | 0 | 42 | 5.41 |
| L_Occipital_lobule | -30 | -84 | -6 | 6.45 |
|  | -36 | -78 | -6 | 5.6 |
|  | -22 | -88 | -8 | 4.78 |
| L_S2 | -50 | -22 | 30 | 6.42 |
|  | -54 | -16 | 18 | 5.65 |
|  | -56 | -14 | 28 | 5.38 |
| R_S2 | 46 | -22 | 20 | 6.42 |
|  | 58 | -16 | 22 | 5.65 |
|  | 44 | -32 | 26 | 5.38 |
| L_Sup_Motor | -6 | 8 | 54 | 4.64 |
| Cerebellum | -22 | -56 | -22 | 5.89 |
| L_insular | -30 | 18 | 10 | 5.22 |
| R_Inf_Front | 34 | 26 | 10 | 5.2 |
| AH (Stimuli vs Baseline) |  |  |  |  |
| R_S1 | 48 | -18 | 50 | 6.93 |
| L_S1 | -46 | -26 | 50 | 4.58 |
|  | -40 | -28 | 50 | 3.47 |
| R_S2 | 58 | -18 | 22 | 6.72 |
|  | 46 | -20 | 20 | 4.33 |
| L_S2 | -48 | -20 | 20 | 6.72 |
| L_M1 | -44 | 0 | 36 | 7.28 |
| R_Occipital_lobule | 32 | -86 | -10 | 4.24 |
| AF (Test vs control) |  |  |  |  |
| R_S2 | 50 | -20 | 24 | 4.13 |
| L_S2 | -48 | -18 | 22 | 5.04 |
| AH (Test vs control) |  |  |  |  |
| R_S2 | 66 | -22 | 22 | 5.94 |
| L_S2 | -58 | -24 | 20 | 5.2 |

### 3.3. Percent signal changes in the somatosensory cortex

We determined individual local ROIs in activated regions identified by the contrast analysis of stimulus versus baseline. Then, we measured the PSC in each ROI and examined its difference between the test and control conditions. We found a significant difference in the contralateral S2 (p = 0.028, Wilcoxon signed rank test) under the AF test and AF control conditions (Fig. 6A). We also found a significant difference in the left S2 (p = 0.0102, Wilcoxon signed rank test) under the AH test and AH control conditions (Fig. 6B). The correlation analysis revealed that the PSC difference between the test and control conditions was not significantly correlated with the PSE difference across subjects in any ROI (Fig. 6C). It suggests that the PSC in the somatosensory cortical regions might not directly elucidate the assimilation effect.

**Fig. 6.**
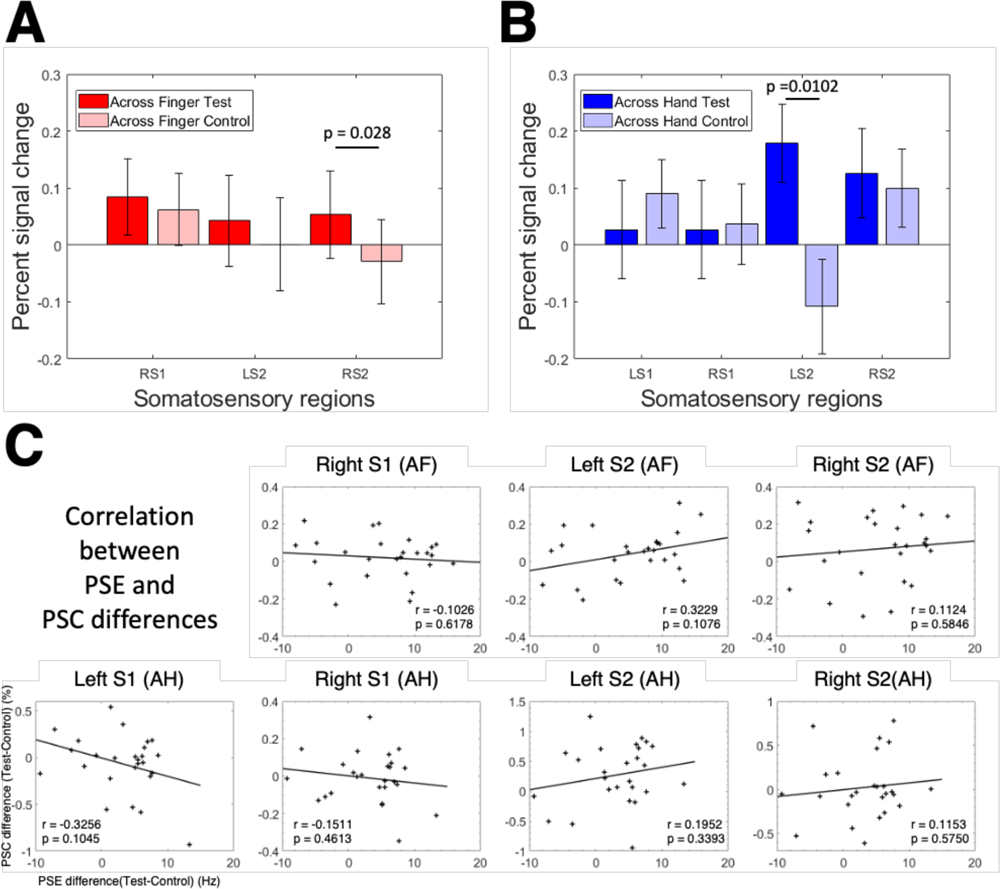
The percent signal changes in the somatosensory regions. (A) The mean percent signal changes (PSCs) in right S1 (RS1), left S2 (LS2), and right S2 (RS2) regions under the test and control across finger (AF) conditions. RS2 shows a significant difference in PSC between the test and control conditions (Wilcoxon signed rank test, p = 0.028). Other regions do not show a significant difference between the test and control conditions. Error bars denote the standard error of the mean. (B) The mean PSCs in left S1 (LS1), RS1, LS2, and RS2 regions under the test and control across hand (AH) conditions. LS2 shows a significant difference in PSC between the test and control conditions (p = 0.0102). Other regions do not show a significant difference. (C) Correlations of PSE differences between test and control with PSC differences between test and control conditions in three regions of interest under the AF condition and four regions of interest under the AH condition. No region shows a significant correlation (ps > 0.05). Each dot denotes data points of individual subjects (N = 26).

### 3.4. Connectivity between somatosensory cortex and other regions

Using the gPPI analysis, we explored the brain regions that increased connectivity with the local ROIs in the somatosensory cortex under the test condition compared to the control condition. In the AF condition, the local ROI in right S1 showed significant connectivity with the medial prefrontal cortex (mPFC) and left inferior parietal lobule (IPL) (FWE corrected p<0.05, cluster size >10, cluster defining threshold p<0.001) (Fig. 7A). However, no significant connectivity was observed regarding other local ROIs in bilateral S2. In the AH condition, the local ROI in left S2 showed significant connectivity with mPFC, right IPL, precuneus, and left temporal sulcus (FWE corrected p<0.05, cluster size >10, cluster defining threshold p<0.001) (Fig. 7B). The local ROIs in left S1, right S1, and right S2 did not show significant changes in connectivity with any other brain regions. These findings suggest that distractor stimuli alter brain connectivity based on the somatosensory region.

**Fig. 7.**
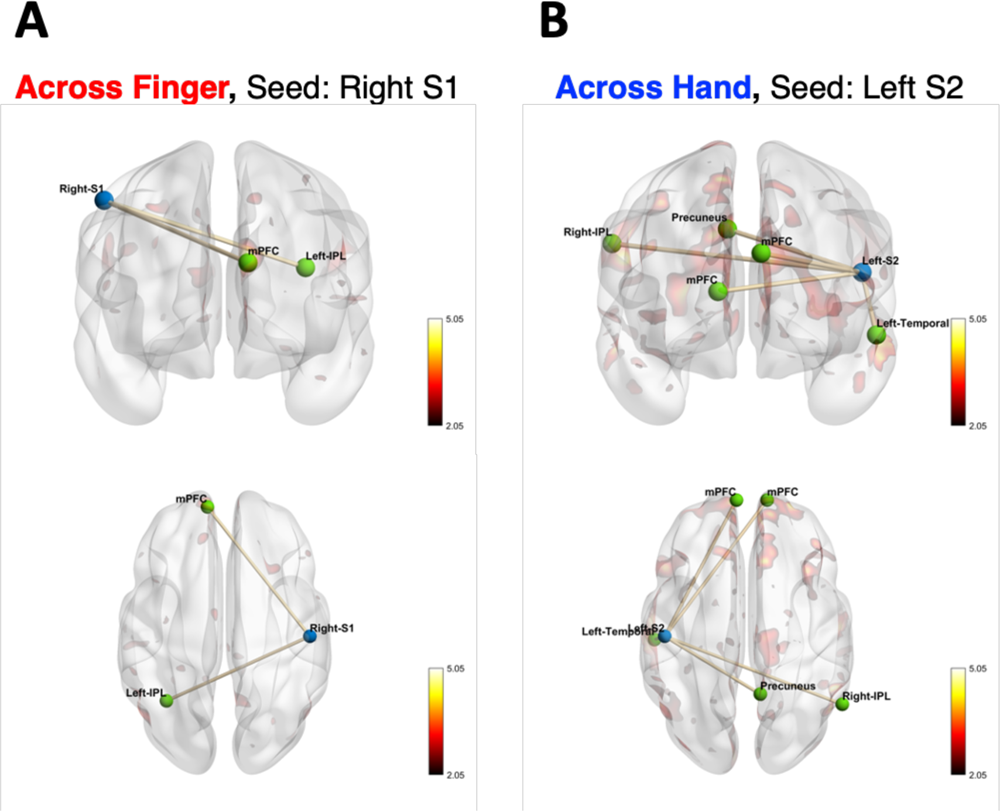
Brain connectivity results of the generalized psycho-physiological interaction (gPPI) analysis. (A) Under the across-finger (AF) condition, the gPPI analysis reveals two significant connections from right S1 (seed region) to the medial prefrontal cortex (mPFC) and left inferior parietal lobule (IPL). No significant connectivity is found when left S2 or right S2 is used as a seed region. (B) Under the across-hand (AH) condition, the gPPI analysis reveals four significant connections from left S2 to mPFC, right IPL, precuneus, and left temporal sulcus. No significant connectivity is found for left S1, right S1, and right S2 as a seed region. Significant connectivity is evaluated with p<0.05 corrected for multiple comparisons at the cluster level, with a height threshold at the voxel level of p<0.001. The blue node indicates the seed ROI, and the green nodes indicate the brain regions that show significant connectivity with the seed ROI. The color maps show the t-values of the gPPI results.

We examined correlations between individual connectivity and individual PSE differences for each inter-regional connection identified by the gPPI analysis above. In the AF condition, individual connectivity between the right S1 and mPFC showed a significant negative correlation with the PSE difference (p = 0.0112, r =-0.4893) (Fig. 8A). In the AH condition, connectivity between the left S2 and right IPL showed a significant positive correlation with the PSE difference (p = 0.0474, r = 0.3923) (Fig. 8B). Other inter-regional connections did not show significant correlations with the PSE difference (p > 0.05). The results of fMRI experiments show that the assimilation effect cannot be explained solely by the response level of the somatosensory region. Instead, it can be attributed to higher cognitive processes, which involve changes in connectivity between the somatosensory region and other brain areas.

**Fig. 8.**
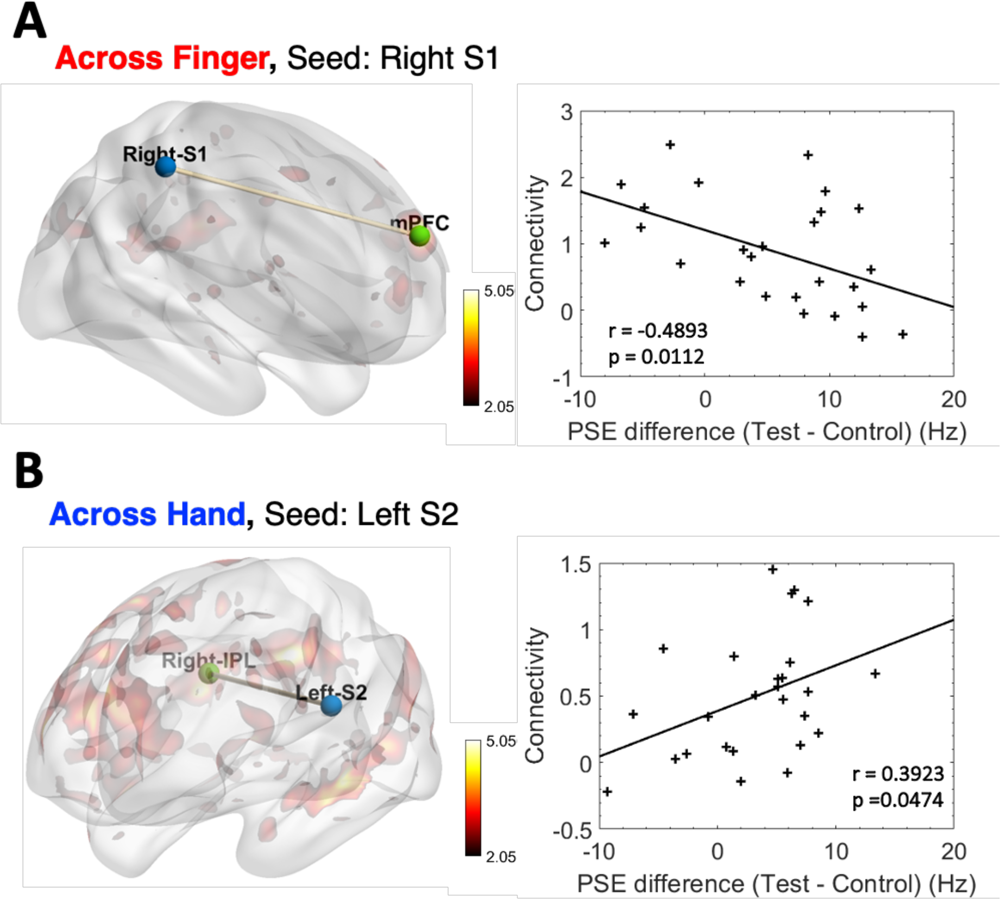
Correlations between individual PSE and brain connectivity. (A) Individual connectivity between right S1 and mPFC shows a significant negative correlation with individual PSEs under the across-finger (AF) condition (p = 0.0112, r = -0.4893). (B) Individual connectivity between left S2 and right IPL shows a significant positive correlation with individual PSEs under the across-hand (AH) condition (p = 0.0474, r = 0.3923). The blue node indicates the seed ROI, and the green nodes indicate the brain regions that show significant connectivity with the seed ROI. The color maps show the t-value of the gPPI analysis results. Each dot denotes individual subjects (N=26).

## 4. Discussion

Considering the parallel processing of tactile afferent signals from the mechanoreceptors at various locations up to S1, it is intriguing to know which stages of neural processing of tactile stimuli in the brain are involved in the occurrence of the assimilation effect. To address it, we examined neural responses involved in the assimilation effect for vibrotactile stimulations. In this study, we found that the right S2 of the AF condition and the left S2 of the AH condition responses were increased by the presence of the distractor stimulus that induced the assimilation effect. However, the PSC in these S2 areas was not correlated with the PSE across subjects, implying that the observed S2 activations might reflect the processing of multiple types of tactile signals together. Also, we found that inter-regional connectivity between the somatosensory cortical regions and other brain regions was increased by the presence of the distractor stimulus. Among these connections, individuals whose perception were more affected by the assimilation effect showed decreased connectivity between right S1 and mPFC in the AF condition and increased connectivity between left S2 and IPL in the AH condition. Therefore, our results suggest that the vibrotactile assimilation effect may be associated with interactions between the somatosensory cortex and mPFC or IPL.

### 4.1. Neural activations in somatosensory cortex

Primary and secondary somatosensory regions showed responses to vibrotactile stimulations to the left or both hands in line with previous studies (Chung et al., 2013; Lamp et al., 2018; Lee et al., 2016). Among these somatosensory cortical regions, the PSC in Right S2 under the AF condition and left S2 under the AH condition was increased when the distractor stimulus was presented together with the target stimulus in the test condition. As S2 is reportedly involved in the integration of somatosensory information (Zhu et al., 2007), these S2 activations may be associated with integrating tactile information of RA-I and RA-II afferents from distinct fingers. However, S2 activations observed in this study may represent cortical responses to high-frequency vibrotactile stimuli (e.g., 200 Hz in this study), as many studies reported more prominent neural responses of S2 to high-frequency vibration than to low-frequency flutter (Chung et al., 2013; Francis et al., 2000; Harrington and Hunter Downs, 2001). Since a 200-Hz tactile stimulus was provided as a distractor in the test condition, although unattended, it might increase S2 responses compared to the control condition under which no high-frequency stimulus was presented. To address this potential compounding, we may need to test whether the contrast of the test against control still activates S2 in an opposite condition where a high-frequency target stimulus (e.g., 200 Hz) and a low-frequency distractor stimulus (e.g., 30 Hz) are presented. Yet, this study could not employ such a stimulation design due to the resonance issue around 200 Hz generated by the PTS used here (see *2.2. Vibrotactile stimuli*). In addition, we observed that individual PSC differences in S2 were not significantly correlated with individual PSE (Fig. 6). Therefore, we drew a conclusion that S2 responses observed in this study may not directly represent the assimilation effect.

### 4.2. In-hand assimilation effect on brain connectivity

We found significant changes in connectivity between right S1 and left IPL and between right S1 and mPFC between the test and control conditions of vibrotactile stimulations on the left hand. Structural and functional connectivity between S1 and IPL regarding tactile perception is well documented (Andersen et al., 1990; Rolls et al., 2023). While regional activities of IPL and S1 did not significantly increased, their connectivity increased when dealing with RA-I and RA-II signals. This result indicates that S1 activity became more related to IPL activity when both the target and distractor stimuli were presented than when only the target stimulus was presented. Yet, no significant correlation between individual connectivity and PSE was found across subjects, suggesting that increased connectivity between S1 and IPL might represent transmitting multiple types of tactile signals.

We also found significant increases in connectivity between right S1 and mPFC. In contrast to S1-IPL connectivity, individual connectivity between S1 and mPFC showed a significant negative correlation with individual PSE. It suggests the engagement of S1-mPFC connectivity in the occurrence of the assimilation effect. Previous research has shown that S1-mPFC connectivity positively correlates with congruency between multisensory stimuli during recognition memory tasks (Van Kesteren et al., 2010). Based on this report, we speculate that when subjects experienced more distraction by the 200-Hz stimulus during the perception of the 30-Hz stimulus in the test condition, the perception of the 30-Hz stimulus to which subjects paid attention might be more dissimilar between the test and control conditions, increasing incongruency between the perception of the 30-Hz stimulus with and without distractor (note that test and control trials were randomly mixed in a block). As such, S1-mPFC connectivity in those subjects would be weaker while the degree of the assimilation effect would increase.

### 4.3. Across-hand assimilation effect on brain connectivity

Across-hand vibrotactile stimulations with target and distractor stimuli changed connectivity between left S2 and other regions, including mPFC, precuneus, left temporal sulcus, and right IPL. These brain regions have been reported to be related to tactile perception. Precuneus is involved in the detection of incongruent tactile stimuli (Kitada et al., 2014). The temporal sulcus reportedly responds to vibrotactile stimulation (Beauchamp et al., 2008; Hegner et al., 2007) and contributes to integrating multisensory information (Renier et al., 2009). IPL plays a role in recognizing tactile stimuli and object shapes (Jäncke et al., 2001) and shows connectivity to S2 during the tactile memory task (Kostopoulos et al., 2007).

However, only individual connectivity between left S2 and right IPL showed a significant positive correlation with individual PSE. Previous studies reported that connectivity between S2 and IPL increased during tactile object recognition (Yu et al., 2018) and vibrotactile stimulation (Chung et al., 2013). IPL is involved in integrating sensory information (Clower et al., 2001). Based on previous findings, those more biased by the assimilation effect might be less likely to suppress the distractor stimulus on the unattended hand, showing increased connectivity between left S2 and right IPL to deal with vibrotactile inputs on both hands. It is noteworthy that the assimilation effect modulated connectivity between S2 and IPL across hemispheres, indicating a possibility that cross-lateral brain connectivity might be engaged in the emergence of the across-hand assimilation effect.

### 4.4. Differences between in-hand and across-hand assimilation effects on brain connectivity

We found a negative correlation of individual connectivity between right S1 and mPFC with individual PSE for the in-hand assimilation effect, whereas a positive correlation of individual connectivity between left S2 and right IPL with individual PSE for the across-hand assimilation effect. Also, we confirmed that individual PSE changes from the control to test conditions did not show a significant correlation between the AF and AH conditions (p = 0.1691, r = -0.2838) (Fig. 9). This result shows that individuals who exhibited stronger assimilation effect under the AF condition did not necessarily show stronger assimilation effect under the AH condition. Furthermore, other studies reported different activation patterns in S1 and S2 when a pair of vibrotactile stimuli were presented to one hand or both hands (Tamè et al., 2014). Taken together, our results imply that the in-hand and across-hand assimilation effects involve different neural circuits.

**Fig. 9.**
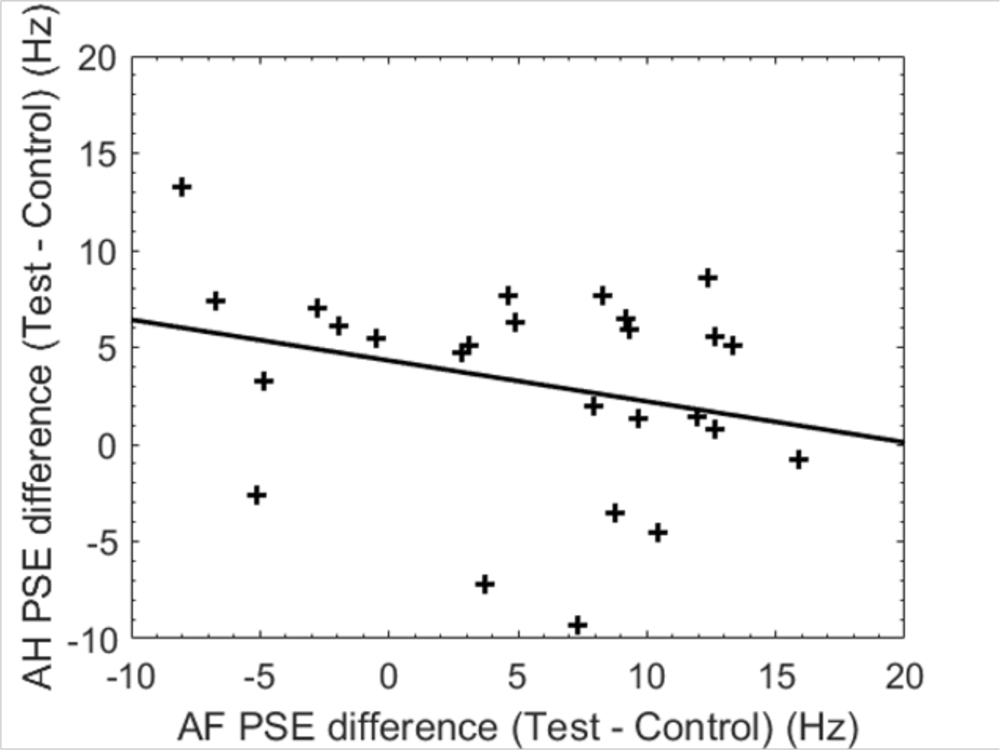
Correlation between across finger (AF) and across hand (AH) PSE differences. The individual PSE differences between the test and control AF conditions do not show a correlation with those between the test and control AH conditions (p = 0.1729, r = -0.2757). Each dot denotes individual subjects (N=26).

### 4.5. Why assimilation, not contrast effects with vibrotactile stimulation?

The assimilation effect describes a perceptual bias towards the distractor stimulus, while the contrast effect refers to a perceptual bias away from the distractor (Kahrimanovic et al., 2009). Among these, the assimilation effect occurred when vibrotactile stimuli with different frequencies were presented. The study by Kahrimanovic *et al*. also reported the assimilation effect by tactile stimulations such that when both the control stimulus and a soft or rough stimulus were presented to the adjacent fingers simultaneously, the subjects showed the assimilation effect of responding to the control stimulus as softer or rougher (Kahrimanovic et al., 2009). Linking to the visual assimilation effect, Kahrimanovic *et al*., assumed that the presence of receptive fields in the somatosensory cortex may play a role in the assimilation by tactile stimulations. If tactile stimuli are presented to the index and middle fingers of which the receptive fields are close to each other, the brain may sum stimuli in the receptive fields and perceptually assimilate the averaged signal.

Likewise, under the AF condition, we can conjecture that vibrotactile afferent signals in the receptive fields of the index and middle fingers might be averaged in S1. Then, S1 transmits this averaged signal to mPFC for tactile discrimination, which leads to the perception of incongruent tactile stimuli and decreased connectivity. Yet, this assumption about the proximity of the receptive fields may not apply to the assimilation effect under the AH condition because it is unknown whether the receptive fields of two different index fingers are close to each other within S2. Thus, the vibrotactile assimilation effect may need to be elucidated by more complex neural processing than a simple mixture of receptive fields.

## Credit authorship contribution statement

**Ji-Hyun Kim:** Conceptualization, Data curation, Formal analysis, Investigation, Methodology, Software, Validation, Visualization, Writing – original draft.

**Dooyoung Jung:** Data curation, Funding acquisition, Formal analysis, Investigation. **Junsuk Kim:** Conceptualization, Formal analysis, Investigation, Methodology, Software, Validation, Writing – review & editing

**Sung-Phil Kim:** Conceptualization, Funding acquisition, Project administration, Supervision, Writing – review & editing.

## Data and code availability

Data will be made available on request.

## Acknowledgments

This research was supported by the Brain Research Program through the National Research Foundation of Korea (NRF) funded by the Ministry of Science, ICT & Future Planning (2022M3C7A1015112) and the National Research Foundation of Korea (NRF) grants funded by the Korea government (MSIT) (No. RS-2023-00302489) and the Alchemist Project (20012355, Fully implantable closed loop Brain to X for voice communication) funded by the Ministry of Trade, Industry & Energy (MOTIE, Korea).

